# Enhanced Detection of Circular Permutations in Protein Databases

**DOI:** 10.1101/2024.08.28.610105

**Authors:** Zundan Ding, Zeji Li, Zitong Zhou, Zeying Wang, Bin Huang, Yue Hu

**Affiliations:** School of Life Sciences, Yunnan Normal University, Kunming 650500, Yunnan, China; College of Bioengineering, Qilu University of Technology (Shandong Academy of Sciences), Jinan 250300, Shandong, China; Kyiv College, Qilu University of Technology (Shandong Academy of Sciences), Jinan 250300, Shandong, China

## Abstract

This study developed a robust method to detect circular permutations in the Protein Data Bank, analyzing 287,081 proteins with sequence lengths under 800 residues. By employing Foldseek and MMseqs2 for similarity searches and refining results with TM-align, icarus, and plmCP, we identified 20,801 potential circular permutation pairs and 3,351 unique circular permutation proteins. These findings have been compiled into **PermuStructDB**, a comprehensive database dedicated to circular permutation proteins. This approach, along with the establishment of **PermuStructDB**, significantly advances our understanding of protein structural variations and evolutionary adaptations, providing a valuable resource for future research in protein engineering and design.

## 1. Introduction

Circular permutation^1^ is a structural rearrangement in proteins where the N-terminal and C-terminal ends are swapped, and the protein sequence is “circularized” by connecting the original termini. This structural variant often retains the same overall fold and function but may exhibit differences in stability, activity, or regulation. These advances highlight the versatility of circular permutation as a tool for understanding protein structure and function, as well as its potential for creating novel biomolecules with practical applications.

Detecting circular permutations in proteins is an important task in bioinformatics and structural biology. Several computational tools have been developed to identify and analyse these rearrangements in protein sequences and structures. These tools^2, 3^, developed by various researchers, provide robust methods for detecting and analysing circular permutations in proteins. They range from sequence-based approaches (like seqCP^4^ and CPred^5^) to structure-based tools (like TMalign_CP^6^) and databases (like Circular Permutation Database) that integrate both types of data. These tools are essential for studying protein evolution, understanding structural diversity, and designing new proteins with desired functions.

The Circular Permutation Database^7^ (CPDB) is a specialized resource dedicated to the study of circular permutations in proteins. This database serves as a crucial tool for the scientific community, providing insights into the structural and functional implications of circular permutations in proteins. The database not only facilitates research in this specialized area but also helps in advancing broader understanding in areas like protein engineering, structural biology, and evolutionary biology. However, the CPDB was published in 2009 and was a derivative database from the Protein Data Bank^8^. At its inception, it contained a curated set of circular permutation data but was limited by its relatively small size and the scope of available structural information at the time. Shortcomings include its static dataset, which may not encompass more recent discoveries or integrate with newer computational methods, potentially limiting its usefulness for current research needs.

Analysing pairwise structural similarities for the entire Protein Data Bank (PDB) presents significant challenges due to the sheer volume of data and computational complexity. With about 300,000 protein structures, performing comparisons for every possible pair is immensely resource-intensive. For example, tools like TM-align, while effective for individual pairwise comparisons, become impractical for large-scale analysis due to their high computational and memory demands. The process requires calculating and storing similarity scores for every pair, which quickly becomes unmanageable as the number of structures increases. Foldseek^9^, on the other hand, is designed to handle large-scale structural comparisons more efficiently, as it can perform comparisons in a more parallelized and optimized manner. However, even Foldseek or MMSeqs2^10^ faces limitations related to processing power and memory, and may require sophisticated infrastructure to handle the massive dataset of the PDB. Both approaches thus struggle with the massive computational load and memory requirements involved in analysing structural similarities across such a vast database.

In this article, we combine these advanced tools to delve into circular permutation in the PDB. Foldseek, integrated with protein language models, facilitates database creation for any protein sequence. It transforms protein sequences into one-dimensional information with structural insights. After screening, we use pairwise structural comparison tools, such as TM-align, icarus^11^, and plmCP^12^, for further filtering and validation. We have simplified the classical process of discovering circular permutations, enabling the identification of circular permutations in the database with a single comparison of two databases (**Figure 1**). Ultimately, we completed a systematic exploration of circular permutations in the PDB up to January 1, 2024, proving the effectiveness of the entire algorithm. The results of this exploration have been compiled into **PermuStructDB**, the latest and most comprehensive circular permutation database, which we are proud to provide to the scientific community. This resource not only validates the robustness of our approach but also serves as a valuable tool for advancing research in protein structural variations and evolutionary adaptations.

**Figure 1.**
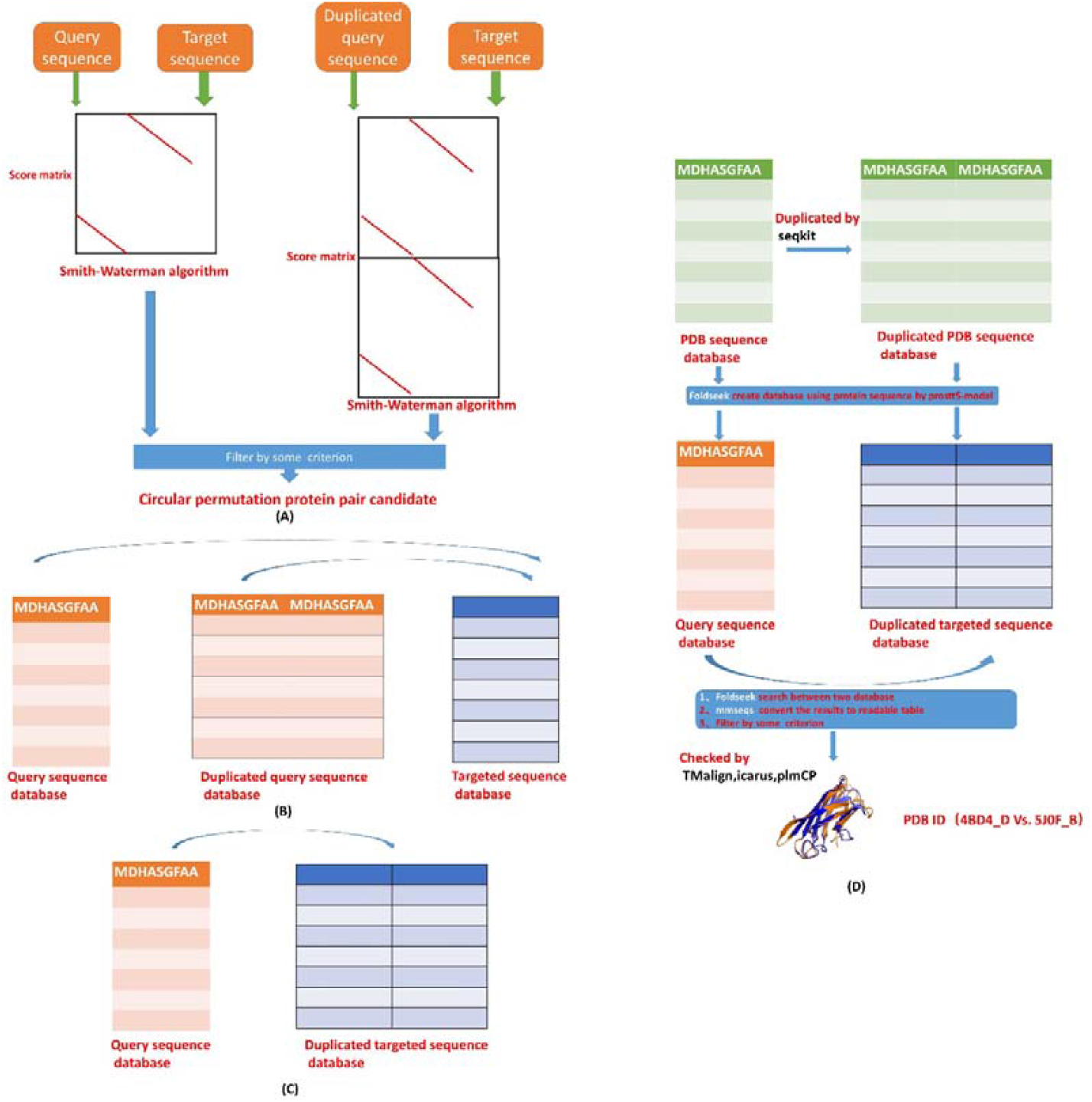
The overall algorithm of database search for the detection of circular permutation: (A)classical circular permutation algorithm;(B) classical circular permutation database search;(C) circular permutation database search used in PermuStructDB;(D) the details of algorithm of the circular permutation detect for PermuStructDB.

## 2. Results

We have obtained 20801 pairs of circular permutation proteins, totalling 3351 proteins. We have made this database (**PermuStructDB**) available on GitHub for public download(https://github.com/YueHuLab/Circular-permutation-in-PDB). Upon further examination of **PermuStructDB**, we first looked at the protein pairs and compared the TM-scores calculated using TM-align without the circular permutation parameter and with the circular permutation parameter (**Figure 2(A)&(B)**). We then plotted probability density plots and scatter plots, which clearly show significant differences between the two and the TM-scores are relatively large, compared to the CPDB database (**Figure 2(E)&(F)**). This indicates that the circular permutation we identified is quite prominent. We also compared the lengths of the circular permutation proteins of **PermuStructDB** with the lengths of all proteins in the PDB database that are shorter than 800 residues (**Figure 2(C)**). We found significant differences between the two, with fewer circular permutation proteins being either much shorter or much longer. The lengths of the circular permutation proteins follow a normal distribution, cantered between 200 and 300 residues. Finally, we examined the proportion of the four CATH^13^ classifications (Mainly Alpha (α): Mainly Beta (β); Alpha-Beta (α/β); Few Secondary Structures:) in the **PermuStructDB** we obtained and compared it with the proportion of the four CATH classifications in the PDB database using histograms (**Figure 2(D)**). Observations and Fisher’s exact test revealed significant differences. Circular permutations are predominantly concentrated in Mainly Alpha (α) and Mainly Beta (β), with very few in the mixed Alpha-Beta (α/β).

**Figure 2.**
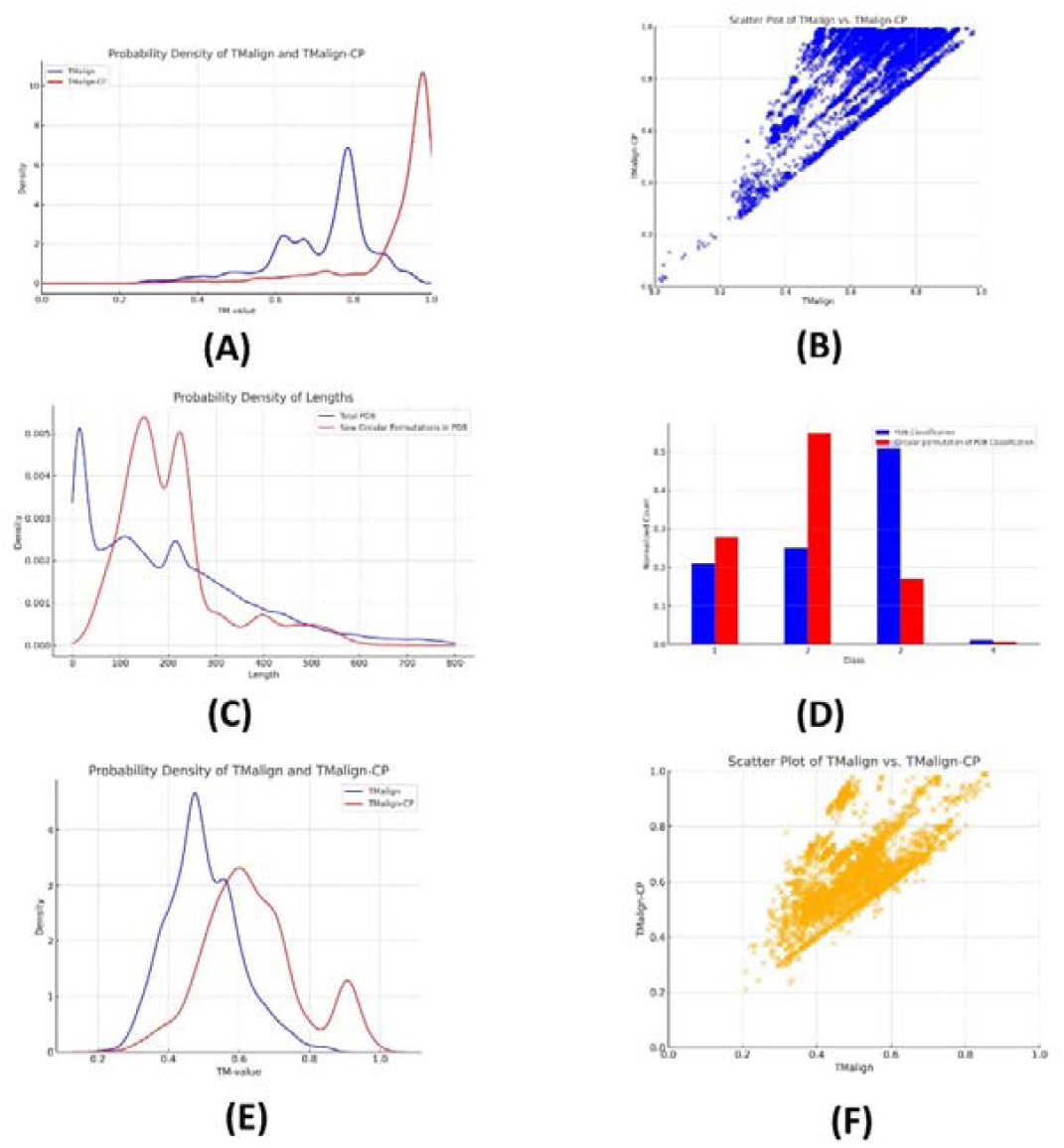
Statistic of the circular permutation proteins:(A) and (B) are the probability density and scatter diagram of the TM-value of TMalign versus TMalign-CP of circular permutation pairs discovered in PermuStructDB respectively. (C) is the length distribution of circular permutation proteins of PermuStructDB and total proteins in PDB;(D) is the histogram of the CATH classification of circular permutation proteins of PermuStructDB and total proteins in PDB; (E) &(F) are are the probability density and scatter diagram of the TM-value of TMalign versus TMalign-CP of CPDB.

We utilized Foldseek in combination with AI models to generate a one-dimensional string representation of protein structures from sequences. This approach is no longer limited to existing protein structures, as it leverages protein language models to effectively provide structural information, ensuring accuracy. We also improved traditional circular permutation methods, simplifying the process of library-to-library searches and completing the task of searching the PDB database. After filtering and further structural alignment, we ultimately identified a substantial number of circular permutation pairs and proteins. These findings were statistically analysed and visualized, laying a solid foundation for discovering circular permutations in the protein universe.

## 3. Discussion

In this study, we have successfully constructed and updated the new PermuStructDB database, which has undergone rigorous testing in terms of both library capacity and quality. A notable departure from previous methods is that we did not employ structure-to-structure searches. Instead, we adopted the Feekseek approach, which converts sequences into structure-embedded strings for alignment and search purposes. This innovative method allows for more efficient and accurate sequence comparisons.

Furthermore, we have optimized the algorithm to streamline the search process, enabling comprehensive comparisons to be completed with a single database query. This optimization significantly enhances the efficiency of the database, making it a more robust and user-friendly tool for researchers.

However, it is important to note that the protein library used in this study is relatively small, comprising approximately 250,000 structures from the PDB database. This choice was made because these proteins have experimentally determined structures, which facilitates a reliable evaluation of the results. Nevertheless, for larger databases, such as the protein universe with hundreds of millions of known sequences, the computational requirements remain substantial. Future advancements in both software and hardware will likely be necessary to fully explore the protein universe.

This database serves as a foundational resource for protein engineering, particularly in the design of circular permutations. It provides a reliable platform for developing tools to assess the feasibility and design of circular permutations. By offering a database grounded in experimental structures, it supports fundamental research on circular permutations and lays the groundwork for further advancements in this field.

Overall, these advancements in the PermuStructDB database not only improve its performance but also provide a more streamlined and effective platform for sequence analysis and comparison, while contributing to the broader exploration of protein engineering and design.

## 4. Method

Inspired by classical circular permutation detection methods (**Figure 1(A)&(B))**, we have developed a more resource-efficient algorithm that requires only a single systematic search (**Figure 1(C)**). The drawback of this method is that it may easily identify identical or similar structures, which can also be viewed as a form of circular permutation, where the N- and C-termini of a protein are connected and then cleaved at similar positions. Fortunately, such cases can be easily checked and filtered based on alignment information (such as positions). After filtering, we use low-throughput but more accurate structural and sequence alignment tools for confirmation.

First, we use Foldseek and MMseqs2^10, 14^ to build and search the database(**Figure 1(D)**).. Starting with the sequences, we use the ProstT5 protein language model module of Foldseek to construct both the query sequence database and the duplicated target database. We then perform searches using Foldseek and convert the results into a readable format with MMseqs2. Foldseek is a fast and accurate tool designed for comparing protein structures. It is particularly useful for large-scale structural comparisons, such as identifying structural homologs in protein structure databases. Foldseek is known for its ability to handle large datasets efficiently while providing high sensitivity in detecting structural similarities. MMseqs2 is a fast, ultra-sensitive tool for sequence searching, clustering, and annotation. It is designed to handle large-scale sequence datasets, offering a balance between speed and sensitivity. MMseqs2 is widely used for tasks that involve searching large sequence databases, clustering sequences, or annotating sequences with functional information. Subsequently, we use the alignment results to filter and select data for further analysis. Next, we use TM-align, Icarus, or plmCP to validate the filtered results. We select TM-align with the CP parameter, and consider those results with an increased TM score as the final results.TM-align is a structural alignment tool that compares the 3D structures of proteins. It is widely recognized for its ability to align protein structures and assess their similarity using the TM-score^15^, which is a measure of structural similarity that accounts for differences in protein size and alignment length. For technology details, please see https://github.com/YueHuLab/Circular-permutation-in-PDB/blob/main/flowchart.txt.

## Data Availability

All relevant data can be found at https://github.com/YueHuLab/Circular-permutation-in-PDB/.

## Code Availability

All relevant code and technology details can be found at https://github.com/YueHuLab/Circular-permutation-in-PDB/blob/main/flowchart.txt.

## Author contributions

Bin Huang and Yue Hu and secured funding and designed the algorithm, who are also guarantors of this work and, as such, had full access to all of the data in the study and takes responsibility for the integrity of the data and the accuracy of the data analysis; Bin Huang, Yue Hu, Zundan Ding, Zeji Li, Zitong Zhou and Zeying Wang performed computation and analyzed the data.

## Competing interests

All authors declare no competing interests.

## Acknowledgements

This work was supported by the National Natural Science Foundation of China (Grant No. 32301061 to B.H.) and was supported by the Yunnan Fundamental Research Projects (Grant No. 202401CF070034 to B.H.) and was supported by Shandong Provincial Natural Science Foundation Committee (Grant No. ZR2016HB54 Y.H.) and was supported by Shandong Provincial Key Laboratory of Microbial Engineering (SME).

## Notes

https://github.com/YueHuLab/Circular-permutation-in-PDB

### Competing Interest Statement

The authors have declared no competing interest.

### Summary of Updates

The details of the article have been refined, and the structure has been adjusted to be more specific.

